# Testing covariance models for MEG source reconstruction of hippocampal activity

**DOI:** 10.1101/2021.04.29.441929

**Authors:** George C. O’Neill, Daniel N. Barry, Tim M. Tierney, Stephanie Mellor, Eleanor A. Maguire, Gareth R. Barnes

## Abstract

Beamforming is one of the most commonly used source reconstruction methods for magneto- and electroencephalography (M/EEG). One underlying assumption, however, is that distant sources are uncorrelated and here we tested whether this is an appropriate model for the human hippocampal data. We revised the Empirical Bayesian Beamfomer (EBB) to accommodate specific a-priori correlated source models. We showed in simulation that we could use model evidence (as approximated by Free Energy) to distinguish between different correlated and uncorrelated source scenarios. Using group MEG data in which the participants performed a hippocampal-dependent task, we explored the possibility that the hippocampus or the cortex or both were correlated in their activity across hemispheres. We found that incorporating a correlated hippocampal source model significantly improved model evidence. Our findings help to explain why, up until now, the majority of MEG-reported hippocampal activity (typically making use of beamformers) has been estimated as unilateral.

## 1) Introduction

Encephalographic functional neuroimaging modalities such as magneto- and electroencephalography (M/EEG) work on the principle of detecting an electromagnetic signature of synchronous neuronal currents in the brain, using sensors on or near the scalp of a participant. The estimation of electrical brain activity given the scalp level measurements is an ill-posed problem and requires additional assumptions or prior beliefs about how the data were generated. One such reconstruction method, beamforming, has enjoyed widespread use by the M/EEG community^1–9^.

Recent studies into the neural dynamics of memory have motivated numerous publications where hippocampal function has been imaged non-invasively. One anomaly is that studies using functional MRI (fMRI) often report bilateral hippocampal activations^10–14^, whereas source-level M/EEG studies typically report unilateral activations^9,15–17^. A common link between many of the M/EEG studies is the reliance on beamformer methods.

Beamforming exploits the temporal covariance of recordings from a fixed array of sensors (such as set of magnetometers or electrodes) and tunes the sensitivity to successive anatomical locations, whilst attenuating all other interfering sources. This data-driven approach makes it a powerful, and noise robust, source reconstruction method, that has been widely used in clinical and cognitive neuroscience^4,18–23^. However, the assumption behind beamformer source reconstruction is that sources are *a priori* uncorrelated. Consequently, there are well-known situations, such as the bilateral evoked response due to binaural auditory stimulation, in which source(s) are mislocalised or suppressed entirely. Here we hypothesise that it is this feature of beamforming that may explain why there is a discrepancy between the M/EEG and fMRI literature on the hippocampus.

Solutions to this correlated source problem have been forthcoming for many years, with a wide variety of approaches published^24–31^. For a detailed history, Kuznetsova and colleagues provide a review of the literature within their own solution to the correlated source problem^31^. One common approach is to collapse two distinct and hypothetically correlated sources into a single source^24,27^, the problem being that as these sources are now correlated *a priori*, and all solutions (regardless of the underlying physiology) will reflect this. Although these solutions are practical in situations where such correlations are known to exist, it is difficult to determine when (or if) such measures are necessary when faced with novel paradigms or data.

In this study, we set out to use commonly available tools and methodology to explore data derived from a task known to engage the hippocampus. We made use of the Empirical Bayesian Beamformer (EBB^32^) to explain competing models of MEG data^33^. The advantage of this formulation is that different priors (or assumptions) can be directly compared in terms of their model evidence (as approximated by Free energy^34,35^). We show how model evidence directly reflected the poorer source reconstruction performance of the conventional EBB in the presence of correlated sources. We then demonstrate how it is possible to use different correlated and uncorrelated priors to directly test physiological hypotheses. We show this in simulation and then in an experimental dataset from a recent beamformer MEG study^16^. We found that models based on correlated hippocampal priors provided a significantly more likely explanation of the MEG data. We also identified an issue specific to the anterior hippocampi, which have highly correlated gain matrices and are therefore difficult to distinguish between using conventional cryogenic arrays.

## 2) Theory

### 2.1) A brief summary of empirical Bayesian source reconstruction

An M/EEG dataset,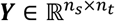 from *n*_*s*_ sensors and sampled over *n*_*t*_ timepoints, generated from *n*_*j*_ current dipoles distributed within the brain, can be described by the generative model

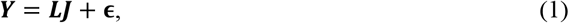

where 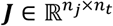 is a matrix which comprising of the activity at each of the sources, 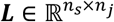 is the gain, or lead field matrix, which contains the dipolar field patterns expected for a single source of for a given location and orientation, and 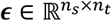 is the unexplained noise, typically modelled as a multivariate Gaussian distribution with zero mean and covariance ***Q***_*ϵ*_. Source reconstruction aims to rearrange Equation 1 such that, given a set of source locations and calculated lead fields, we can obtain ***J***. Here we approached the problem within in a Bayesian, maximum *a posteriori* framework^33,36–38^. Assuming ***J*** is a multivariate Gaussian process, with zero mean and covariance ***Q***_*j*_, the source estimates can be expressed through Bayes’ theorem as

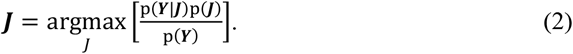

in other words, the distribution of sources which maximises the posterior p(***J***|***Y***). The likelihood and prior are assumed to be multivariate Gaussian processes with zero-mean, such that,

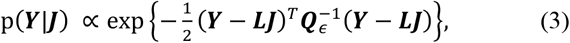

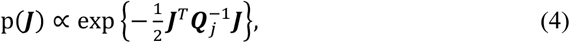

whilst p(***Y***) is assumed to be constant. The derivation of Equation 2 has been covered previously^38,39^, but the end result is a source estimate of the form

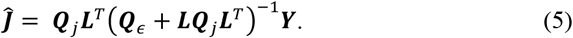

In this expression the lead fields are analytically solved based on physical models of the head^40–43^. The noise covariance matrix 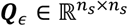 can be directly measured from an empty room recording, but is often assumed that noise from the sensors is independent and individually distributed, such that 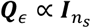 where 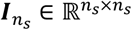 is an identity matrix. The source covariance matrix 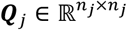 contains the priors or assumptions that differentiate source reconstruction methods from one other^44^. For example, the *minimum norm* solution (MNE^45^) assumes that all sources are considered independent and equiprobable (i.i.d.), such that 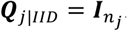 EBB^32^ also assumes independent sources (off diagonal values of ***Q***_*j*|*EBB*_ are set to 0) but each diagonal element corresponds to the (beamformer estimated) variance of a single source. For a given source at a location/orientation *θ*

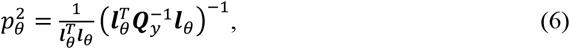

where 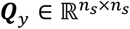 is the sensor-level covariance matrix and 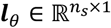 is the corresponding lead field vector. Defining the vector 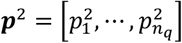, we finally generate the source covariance prior for beamforming

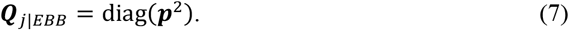

### 2.2) The correlated source Empirical Bayesian Beamformer (cEBB)

Beamformers are known to perform poorly when the underlying current distribution contains distal correlated sources^6,24,25,46^. Put simply, the signal from the partner source at a different location is seen as noise and suppressed. To counter this in our prior construction, we modified the lead field vector in Equation 6 to define a source space where each dipole of interest is paired to another, each pair mapping to a single lead-field^24^. Here the modified variance term, *p*′^2^ for location *θ* is

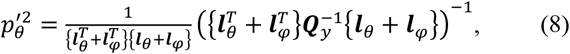

where *φ* is the location/orientation of the second source which correlates with the source at *θ*. Likewise, the variance at *φ* would be the same, i.e. 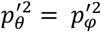 Consequentially, this requires prior knowledge of where this second source is expected to be, but typically it is assumed to be the homolog in the contralateral hemisphere. We also note that not every source need to necessarily to be correlated with another, only w regions of interest if required. By concatenating all the variance measurements into one vector, ***p***′^2^, we generated our final covariance matrix

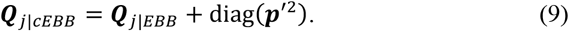

We note that technically we are still assuming independence between all sources, but here we adjust the sensitivity to sources in the prior that we would have otherwise missed with the standard EBB. Note also that the solution is the sum of the original EBB solution and the solution composed of correlated pairs. We briefly investigate the effects of using exclusively one formulation or the other in the Supplementary Information.

## 3) Methods

### 3.1) Simulations

We simulated MEG datasets based on three different scenarios in two different regions of the brain (one cortical region, Heschl’s gyri and one subcortical, the hippocampal regions):

1. A single source generating a 20 Hz sinusoid of amplitude 10 nAm, located in the right hemisphere.
2. Two uncorrelated sources, one a 20 Hz sinusoid of amplitude 10 nAm, located in the right hemisphere. The second source being a 10 Hz sinusoidal source of amplitude 10 nAm, located the homologous region in the left hemisphere.
3. Two correlated, homologous sources in the left and right hemispheres, consisting of two 20 Hz sinusoids (of the same phase) at 10 nAm.

For illustrative purposes, a cartoon depiction of the simulations can be found later in Figure 1A. In each of the three cases, sources were active for 1 second with a 6 second inter-trial-interval, for 10 trials. All simulations were based on a 275-channel SQUID MEG system (CTF, Coquitlam, BC) set in a 3^rd^ order synthetic gradiometer configuration. The locations of three fiducial coil positions (nasion, left and right preauricular) relative to the sensors were derived from a previous experiment^16^. We used canonical rather than individual MRIs for source reconstruction^35,47^. The source space was based on the template anatomical provided with SPM12 (https://www.fil.ion.ucl.ac.uk/spm), with cortical surfaces extracted from the boundary between the pial and white matter. These surfaces were downsampled, such that the 4096 vertices per hemisphere corresponded to the locations and the mesh vertex normals corresponded to the orientations of the dipoles in our source space (used for both the simulations and source reconstruction of the resultant data). In addition, a mesh corresponding to the hippocampus was incorporated in the source space^48^. The source space was then affined transformed (using 7 degrees of freedom) such that the anatomical fiducial locations of the template MRI were aligned to the locations of the fiducial coils relative the MEG sensor array. The lead fields for each of the other sources were generated using a single shell model^41^, where the conductive model geometry was based on a convex hull mesh of the brain/CSF boundary, derived from the transformed template MRI. The SNR of simulations was varied, from 0 dB to -40 dB in steps of 5 dB, sensor noise was sampled from a Gaussian distribution. Data were simulated within SPM12.

**Figure 1:**
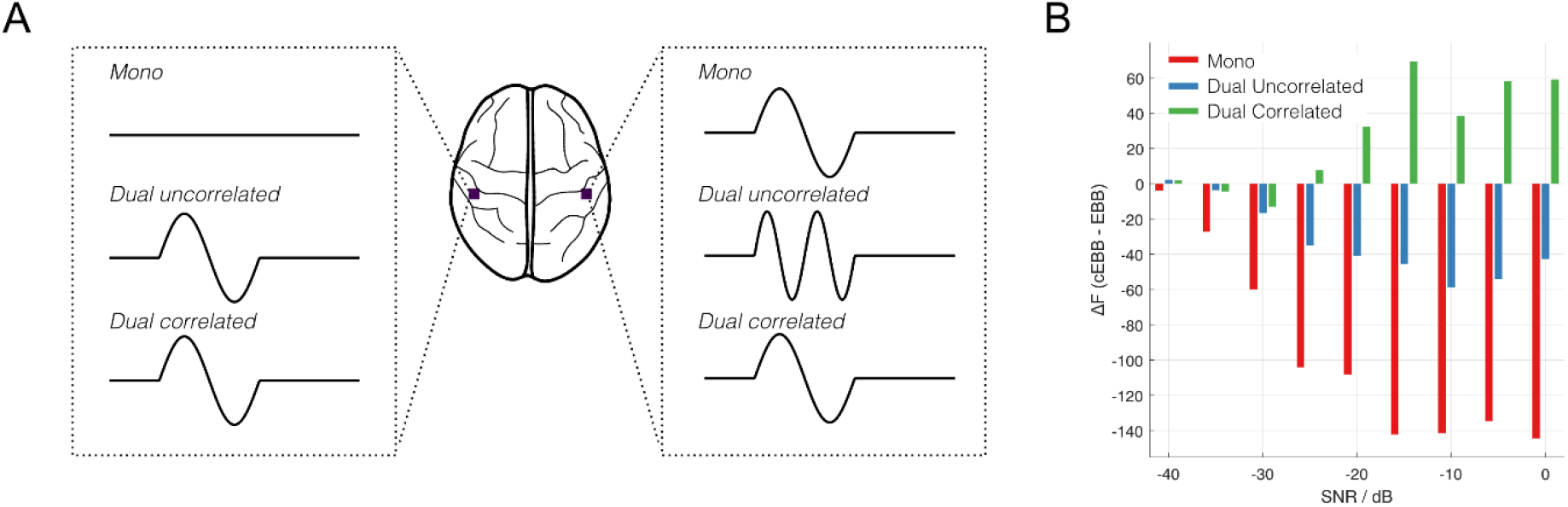
Depiction and model evidence results of the simulations comparing Empirical Bayesian Beamforming (EBB) to correlated EBB (cEBB) inversions. A) Cartoon diagram explaining the simulation types, in this case two sources placed in Heschl’s gyri were set to be either unilateral, uncorrelated or correlated bilateral. B) Model evidence comparison between the correlated and uncorrelated beamformer priors for sources simulated in Heschl’s gyri. Positive values in model evidence indicate that correlated priors (cEEB) were more likely models compared to uncorrelated ones (EBB).

### 3.2) Experimental data

Twenty two native English speakers (14 female, aged 27±7 [mean±SD] years) participated in a previously published study^16^ that involved generating novel scene imagery, a task known to engage the hippocampus. The study was approved by the University College London Research Ethics Committee and all participants gave written informed consent.

The detailed paradigm and experimental considerations can be found in elsewhere^16^ but a summary follows. On any one trial, participants were asked to either imagine a scene, a single isolated object floating against a white background, or count in threes from a specified number. The stimulus type (either, “*scene”*, “*object”* or “*counting”*) was delivered via MEG-compatible earphones (3M, Saint Paul, MN). The participant closed their eyes and awaited the auditory cue of the scene (e.g. “*jungle”*) or object (e.g. “*bottle*”) to imagine, or the number to count in threes from (e.g. “*sixty*”). The participant then had 3000 ms to imagine or count until a beep indicated the trial was over. The participant then opened their eyes. Seventy-five trials of each condition were presented in a pseudorandom order. Note for this investigation, we used all conditions together (i.e. no contrasts) for our analyses.

Data were collected using a 275-channel MEG system (CTF, Coquitlam, BC) at a 1200 Hz sample rate, with 3^rd^ order synthetic gradiometry applied. All participants wore three head position coils, placed on the nasion and left/right preauricular points of the head. These coils were energised prior and after each recording block to establish the locations of the sensors relative to these fiducial points. Trials were visually inspected for SQUID resets and extensive electromyographic artefacts, with trials containing either omitted from further analysis.

### 3.3) Source reconstruction

All source reconstruction was undertaken within SPM12’s DAiSS toolbox (https://github.com/spm/DAiSS). The source space was again defined using a template MRI and its associate cortex/hippocampal mesh as described in Section 3.1. Again, dipole locations were the defined as the mesh vertices and orientation was set to the vertex normal of the mesh. The data covariance matrix ***Q***_*y*_, was calculated using discrete-cosine-transform to filter the data, and decomposed into 4 temporal modes per trial prior to covariance calculation. For the simulations, data were filtered to the 1-48 Hz band. For the experimental data we selected the theta (4-8 Hz) band, as that has been frequently implicated in previous electrophysiological studies of the hippocampus^9,16,49–53^.

The empirical Bayesian scheme involves the optimal weighting of the priors defined in the source covariance matrix ***Q***_*j*_ and an i.i.d. noise matrix ***Q***_*ϵ*_, to the original data covariance:

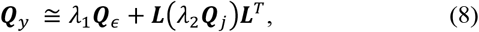

Where *λ*_1_ and *λ*_2_ are non-negative hyperparameters that are estimated by maximising Free energy, *F*, (a lower bound for the model evidence^33^) using a Variational Laplacian estimator^38^, such that

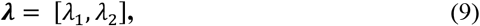

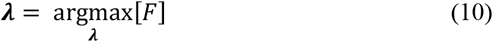

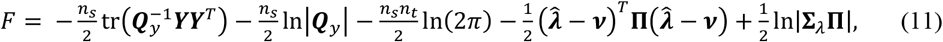

where the prior and posterior distributions *λ, q*(*λ*) and *p*(*λ*) are assumed to be multivariate Gaussians, such that

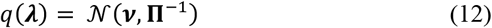

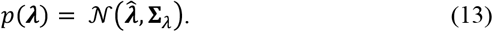

The mean ***ν*** and precision Π of the priors are chosen to be uninformative, such that the *q* is a distribution with near-zero mean (*e*^-4^) and low precision 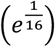 to cover a wide parameter space. Note Π is also a scaled identity matrix. The final optimized free energy (or model evidence) allows us to compare different models (i.e. different formulations of ***Q***_*j*_) of how the data were generated.

In the case of cEBB, we needed to specify different forms of ***Q***_*j*_ depending on which sources were correlated. One such prior was that all homologous source pairs in the two hemispheres were correlated, an assumption made when reconstructing the simulated data. For the experimental data, we also looked at priors containing correlated sources only in the hippocampi but not the cortex and vice versa. For locations where we did not expect correlated sources, we used the definition of source variance in equation 6 rather than equation 8-9 when constructing the prior. From here a source weights matrix 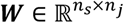 for all sources can be generated using

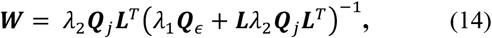

where each column represents a single source. If a source at location *θ* has a corresponding source weights vector 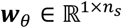, then the reconstructed source power is

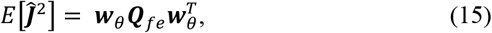

where 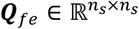 is the data covariance matrix filtered and epoched in to a sub-band and time window of choice.

In the empirical group analysis, we computed the power estimate (equation 15) across the cortical and hippocampal meshes. All power estimates were reconstructed onto a canonical mesh (transformed differently depending on the locations of the fiducial coils within the MEG sensor array) with source spacing of 5 mm on average. This means that vertex N for each participant corresponds to approximately the same location in MNI space. Glass-brain plots were used to depict this mesh in MNI coordinate space. These estimates were then smoothed (along the mesh) using a Green function as the smoothing operator^32^, which gives a local coherence of about 10 mm. We choose 10mm to allow for some intra-subject variability and to increase the sensitivity of the subsequent group statistics. We compared the EBB to various cEBB solutions through a paired t-test of the difference between these smoothed (surface based) power estimates. False Discovery Rate (FDR), as implemented in SPM12, was used to control the false positive rate.

### 3.4) Bayesian Model Comparison

For the simulations that consist of a single dataset, the best model is the one that has the highest model evidence. However, for group studies we may have one candidate model that shows large but random changes in model evidence across the group, or another model that displays modest but consistent improvements. Here, we employed a random effects analysis^54,55^ to inform us which model was more likely or more *frequent* to prevail in a population. From the analysis, two omnibus statistics are useful for our purposes; first, if we have two competing models, m_1_ and m_2_ based on our group-level data G, we can calculate the protected exceedance probability (PXP; ϕ) that one model would be picked over the other:

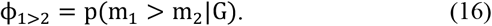

Within the calculation of ϕ, we also quantify the Bayesian Omnibus Risk (BOR) defined as the posterior probability that model frequencies are equal^54^. This could be considered analogous to the classical p-value in frequentist statistics such that BOR<0.05 means that it is unlikely that the model frequencies are the same.

### 3.5) Data Availability

The EBB scheme with options for correlated sources is available within DAiSS toolbox, with an implementation directly within SPM12 arriving in a future update. The full pipeline to run and analyse the simulations is available at https://github.com/georgeoneill/EBBcorr. The experimental MEG data are available upon reasonable request – please contact EAM.

## 4) Results

We divide the results into three main sections. The first section demonstrates the basic principal of operation and the metrics we wished to use in a set of shallow source simulations; the simulation scenarios are familiar and have been explored in many other studies. In the second section, we specifically examined sources on the hippocampal envelope. We had not anticipated this, but for sources located at the anterior hippocampi, the limitations of MEG system sensitivity interacted with the inference we wished to make. Finally, we applied the machinery to data from empirical recordings designed to engage the hippocampus.

### 4.1) Shallow source simulations

We first examined three possible scenarios within Heschl’s gyri and nine different SNR levels in simulation. Figure 1B shows the differences in model evidence (Δ*F*) for cEBB relative to EBB inverse models. Negative values indicate that the EBB solution was more likely than the cEBB. For a single simulated source (Mono; red bars in Fig. 1B) we observed that the cEBB (correlated priors) model was a much less likely description of the data. These differences in model evidence are relatively constant between SNR values of -15 dB to 0 dB (Δ*F* ≈ -140), beyond which we see a reduction in the magnitude of differences as SNR worsens. Similarly, for two uncorrelated sources (blue bars in Fig. 1B) the EBB model provided a more likely description of the data (Δ*F*= -46 at -15 dB). However, in the case of two correlated sources (green bars in Fig. 1B), we noted an increase in model evidence when considering correlated source priors (Δ*F*=69 at -15 dB). We also observed a failure to identify the correlated sources at SNRs lower than -25 dB.

Figure 2 shows the spatial distributions of the source priors for three source inversions. The priors used were EBB, cEBB and, for comparison, an additional vanilla IID model (in which the source prior consisted of an identity matrix) is also shown. Again, these were based on the simulations in Heschl’s gyri (with an SNR=-10 dB) for illustrative purposes. The IID model was not data dependent and so every source location had the same source variance *a priori*, whereas the beamformer models showed differing data-dependent source variance maps. For the mono simulations we noted that the EBB prior highlights only one source, whereas the cEBB prior projected that single source into both hemispheres. This projection can be faintly seen in Figure 2 (marked with a dashed circle), but as the cEBB prior also included information from the EBB prior (it was the sum of the standard and modified priors) this effect was reduced. We investigate the effect of removing the standard EBB prior (from the sum) in the Supplementary Information (see Supplementary Figures 2-3). The priors for the dual (uncorrelated) simulations gave priors that were topographically similar, whilst for the correlated sources, the priors’ topographies diverged. EBB estimated the largest source variance to be in the centre of the brain (and in medial areas of cortex) rather in either Heschl’s gyrus. Conversely, the cEBB prior estimated a variance distribution peaking around the locations of the simulated sources.

**Figure 2:**
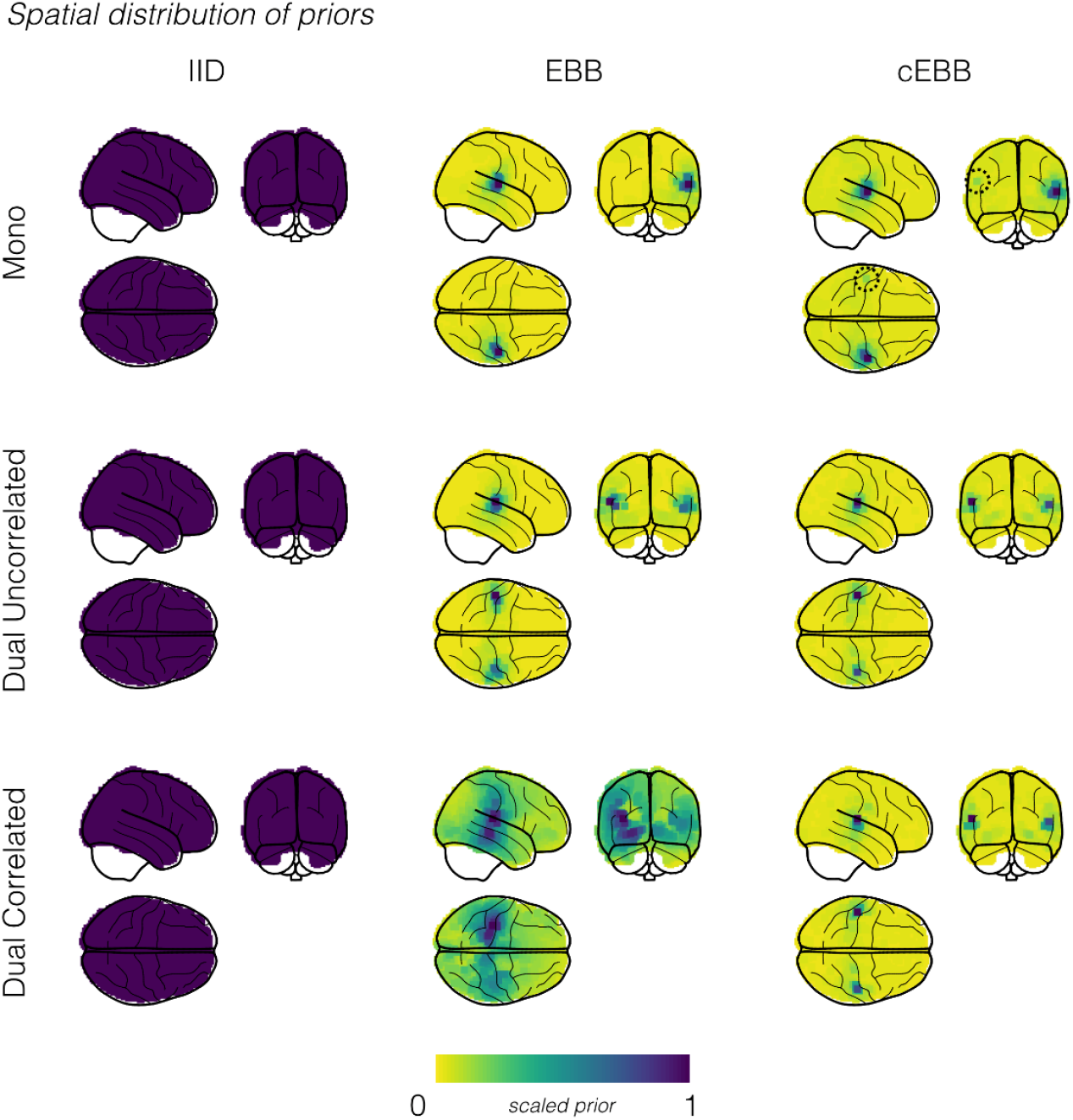
Spatial distributions of the source priors and reconstructed source power plotted onto glass brains for simulations within Heschl’s gyri. Results have been transformed into MNI space. A) The spatial distribution of the source priors, scaled relative to their maximum value for three inverse models, Minimum Norm Estimation (IID), Empirical Bayesian Beamforming (EBB) and correlated source Empirical Bayesian Beamforming (cEBB). The dashed circle highlights a case where cEBB will estimate variance of a non-existent source in the contralateral hemisphere due to the presence of a genuine source in the ipsilateral hemisphere.

Figure 3 shows glass brain plots of oscillatory power estimates between 8-22 Hz (recall simulated sources were generated at either 10 or 20 Hz) during the stimulation period (0-1000 ms) for the three inverse models and the three simulation types at -10 dB. The left most column, which has the power maps for the IID model, show that it localised the source(s), with power biased towards the most superficial cortical locations; a characteristic of unnormalised MNE source reconstructions^56,57^. The beamformer models for the non-correlated source simulations localised power with spatially similar profiles to each other, with their maximal power in each hemisphere close to their simulated locations (IID mean location error (MLE): 13.4 mm; EBB and cEBB MLE: 3.1 mm). Note that despite cEBB estimating variance in the contralateral hemisphere within its prior, the source was predominantly localised in the (correct) ipsilateral hemisphere. For the correlated source simulations, we observed that the EBB model resulted in a quite different topography compared to cEBB; the power distribution more closely resembled the MNE power map for a correlated source model. cEBB in comparison, correctly localised both sources with less localisation error (EBB mean localisation error: 13.8 mm; cEBB mean localisation error: 3.8 mm). In summary, there are three main points to note in this figure. The IID was robust to the correlation structure in the data but tended to produce superficial source estimates. The EBB algorithm produced precise source estimates for uncorrelated sources but in the case of correlated sources failed gracefully, producing an IID-type solution. Finally, the cEBB produced an accurate solution for the correlated scenario. It is interesting to note that the cEBB solution is asymmetric (Figure 3) whereas the prior is symmetric (Figure 2 and Supplementary Figure S3) This highlights one the key differences between the EBB and standard beamformer solutions (which would also be symmetric). Here we develop the prior ***Q***_*j*_ based on beamformer assumptions but the general solution (equation 14) is also determined by the data and the standard lead field matrix (with no additional covariance structure).

**Figure 3:**
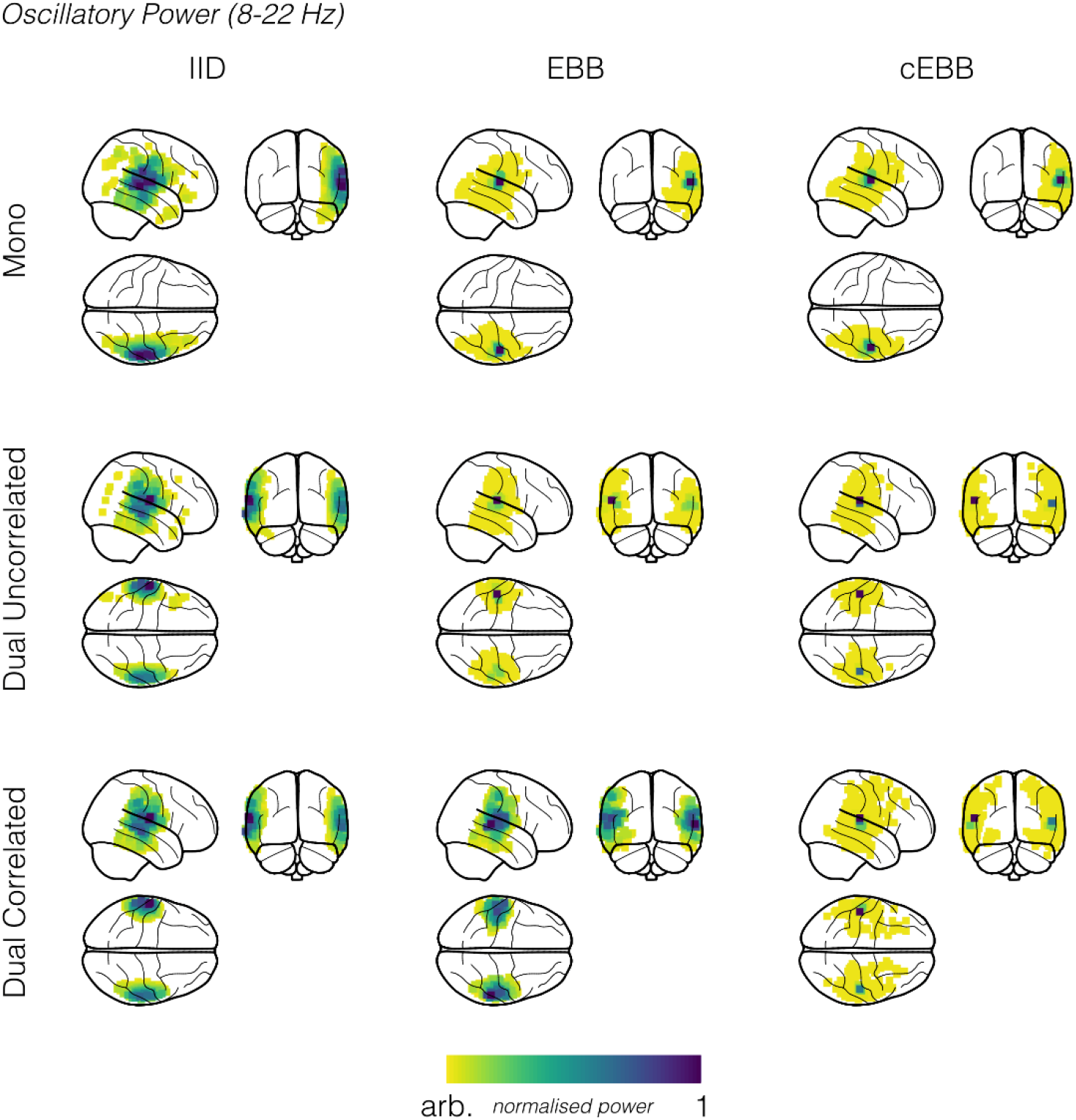
Spatial distributions of the reconstructed source power for unilateral, dual-uncorrelated and dual-correlated simulation scenarios (SNR = -10 dB). When sources were uncorrelated, EBB and cEBB inversion models gave similar results, whereas in the presence of correlated sources, EBB power fell back onto a topography akin to IID. Although qualitatively cEBB reconstructs all three scenarios it is only objectively the best model (see model evidence scores in figure 3) for the dual-correlated sources. Results have been transformed into MNI space for visualisation purposes.

### 4.2) Hippocampal source simulations

The results of the hippocampal simulations are showing in Figure 4. Panel A shows the model evidence from all three simulation scenarios across the 9 SNR levels for a pair of sources in the posterior of the hippocampal body (left plot) and the anterior hippocampus (right plot). In the hippocampal body, we observed very similar behaviour to that found in Hechl’s gyri. It correctly identified that the EBB was a more plausible model for single sources and the case of dual uncorrelated sources, with cEBB being awarded with better model evidence scores for the dual correlated sources. However, for sources in the anterior hippocampus (Figure 4A, right panel), whilst there was a clear distinction between models when the underlying source was unilateral rather than bilateral, sources that were uncorrelated in simulation were more likely (Δ*F* ≈ 10 for most SNR levels) under cEBB prior (i.e. our inference would suggest they were correlated). Figure 4B shows the spatial distribution of model evidence changes when comparing EBB to cEBB for dual uncorrelated sources with an SNR of -10 dB. We observed that for the majority of the hippocampal sources the model evidence suggested that uncorrelated priors were (correctly) more likely, but toward the anterior we noted a transition to where the correlated model was incorrectly classified as more plausible. Figure 4C shows a binarised representation of the model evidence scores; blue areas are where EBB was correctly identified as the winning model, and the red areas where cEBB was favoured. This red zone, in which uncorrelated sources appeared as correlated, was exclusively in the anterior hippocampus. Figure 4D is a plot of the correlation of the lead fields between homologous source pairs in the hippocampus. In the anterior hippocampus, the lead fields between a source and its homolog were highly (r > 0.7) correlated (i.e. occupied a very similar portion of MEG signal space).

**Figure 4:**
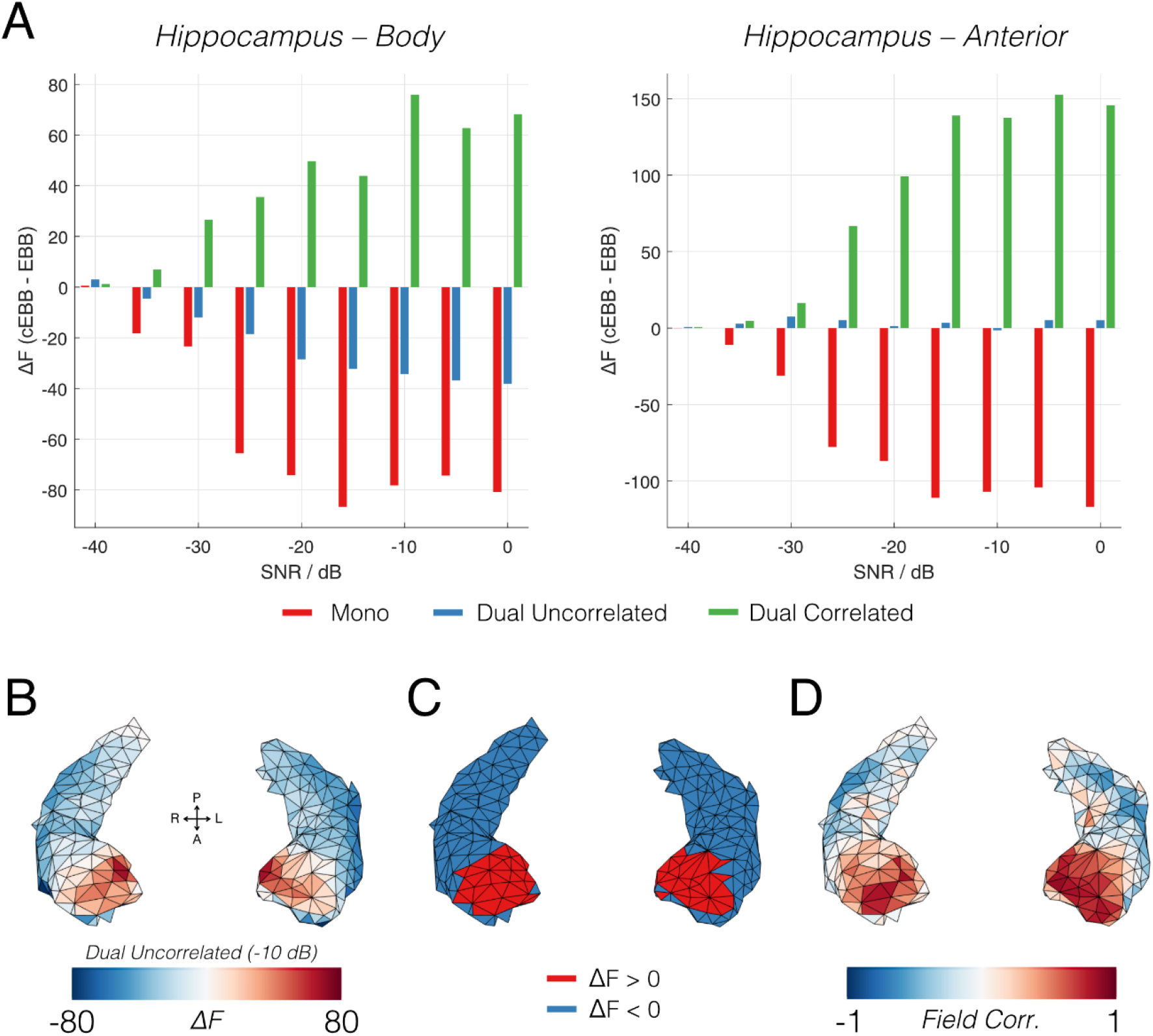
Results of simulations within the hippocampus. A) Model evidence scores of the three simulation scenarios across nine SNR steps for sources in the posterior of the hippocampal body (left) and anterior hippocampal (right). B) Spatial map of the difference in evidence between cEBB to EBB models (cEBB-EBB) for a pair of uncorrelated sources with an SNR of -10 dB. The changes were negative in most cases (i.e. an uncorrelated EBB was a more likely model) but in the anterior we observed that the correlated model was erroneously selected to be more likely. C) A binarised map to show where the simulations correctly identified the underlying data was uncorrelated (blue), and incorrectly suggested to be correlated (red). D) A map of the correlation of the lead fields between homologous source pairs in the hippocampi. Note the strong positive correlation between the anterior hippocampal pairs.

### 4.3) Experimental data

In the previous sections we showed how model evidence can be used to select between different source covariance priors. The question we considered next was how best to describe data originating from the human hippocampus. We reanalysed data from Barry et al.^16^ using four different models: standard EBB using exclusively uncorrelated sources; cEBB with homologous correlated source pairs for the whole brain (across both hippocampi and cortex); cEBB with homologous pairs of correlated sources in the hippocampi only; cEBB with homologous pairs of correlated sources in the cortex but not hippocampi. Figure 5A shows the comparison between competing generative models used to describe data in the 4-8 Hz band in the 0-3000 ms window during all valid trials (across all conditions) for the cohort of 22 participants^16^. Positive values indicate a model is a more likely description of the data than EBB. The first model, where correlated sources could exist in either the cortex or the hippocampus, displayed the largest changes in model evidence compared to EBB (red bars; Δ*F*= 15 ± 32 [mean ± SD] across subjects) followed by restricting the correlated sources to the cortex (blue bars; Δ*F*= 13 ± 30), and the smallest changes by restricting correlations to just the hippocampi (green bars; Δ*F*= 5 ± 3). However, as is evident in Fig. 5A, not all the changes were positive. For the models that contained correlated cortical sources, 5/22 participants exhibited a reduction in model evidence, whereas for the model with the correlated sources existing in only the hippocampi, we observed increases in model evidence in all 22 participants.

**Figure 5:**
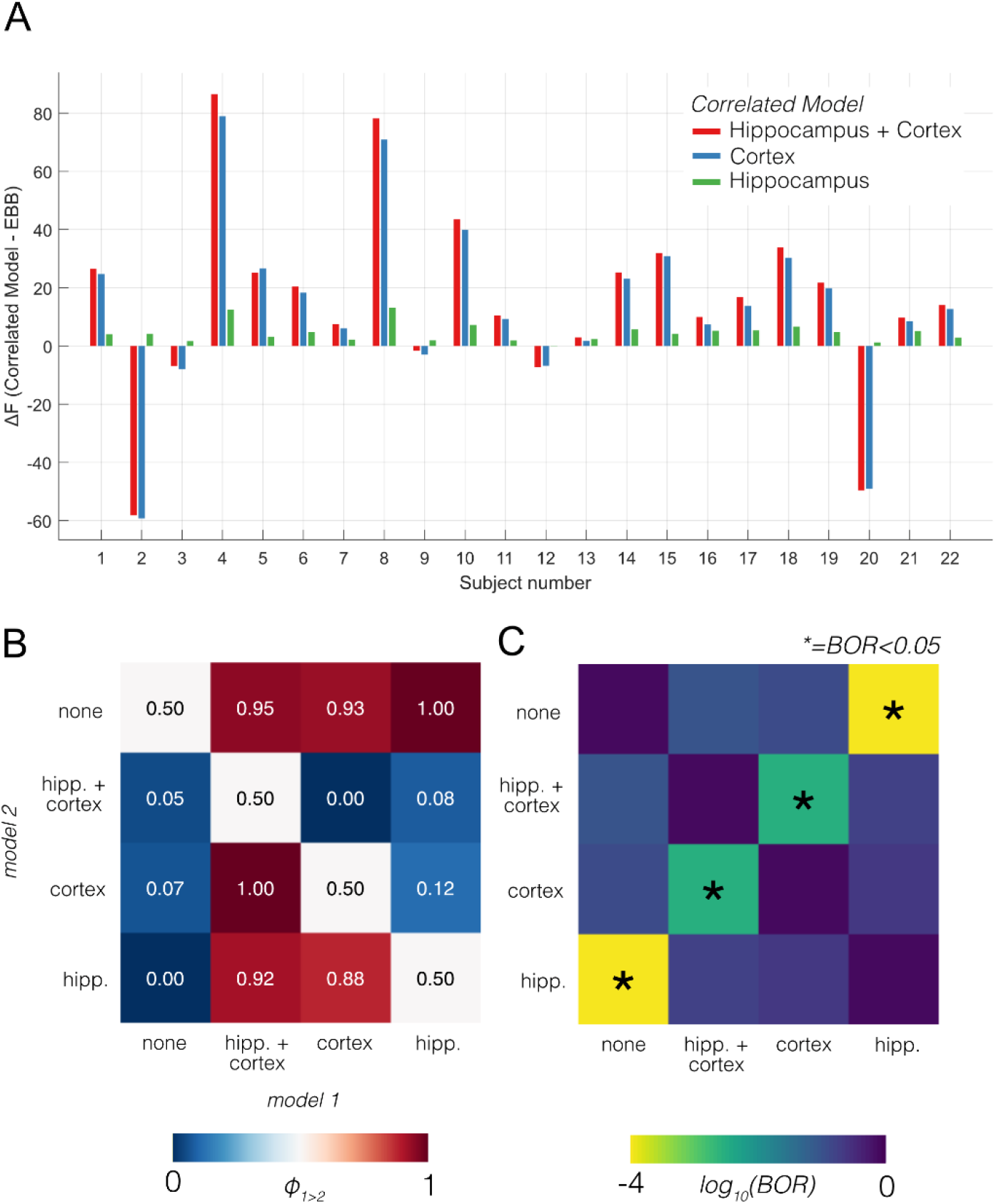
Evidence for four generative models of the experimental data set. A) The changes in model evidence compared to EBB when differing elements of the model were allowed to be correlated: either all sources (red), only the cortical sources (blue) or hippocampal sources (green). B) A random effects pairwise comparison of models in terms of protected exceedance probabilities (PXP; ϕ). High values (red) indicated model 1 was more likely. C) Bayesian Omnibus Risk (BOR), which quantifies how likely it is that a pair of models have the same frequency of occurrence across a group. Asterisks mark comparisons in which model frequencies were clearly different (BOR < 0.05). Note the colour-scale of the BOR values are on a log10 rather than linear scale.

The next step was to determine the most likely model (or most frequently chosen model across the population) using Bayesian model selection^54^. Panel 5B shows the results of the pairwise exceedance probabilities, ϕ between all competing model combinations. Values above 0.5 (red squares) indicate that models labelled along the column are more likely than the model labelled along the row. We found that all models with correlated sources were more likely than the uncorrelated source model (top row, ϕ > 0.9) but, importantly, if the difference between two models was only that the hippocampi were correlated in one and not in the other, then the former model was most frequently chosen in all cases (ϕ = 1). Figure 5C shows the Bayesian Omnibus Risk (BOR), the probability that there is no difference in how often a model would be chosen from across a population. All models involving correlated hippocampal sources were deemed distinct (BOR<0.01) from those without.

Figure 6 shows the group-level FDR-corrected paired-T-contrasts comparing theta band power (4-8 Hz) between source estimates based on uncorrelated (EBB) source models and correlated models containing the hippocampus. Contrasting the correlated hippocampal source model to standard EBB (Fig. 6, left), we noted a bilateral increase in theta power reconstructed in the anterior hippocampus. The right column of Figure 6 shows the contrast between EBB and the cortex-plus-hippocampus correlated model, where we also observed significant changes in power (p < 0.05; whole brain FDR corrected) in the anterior hippocampal regions and anterior/polar temporal cortices. We also observed significant clusters of increased reconstructed power in the occipital cortex.

**Figure 6.**
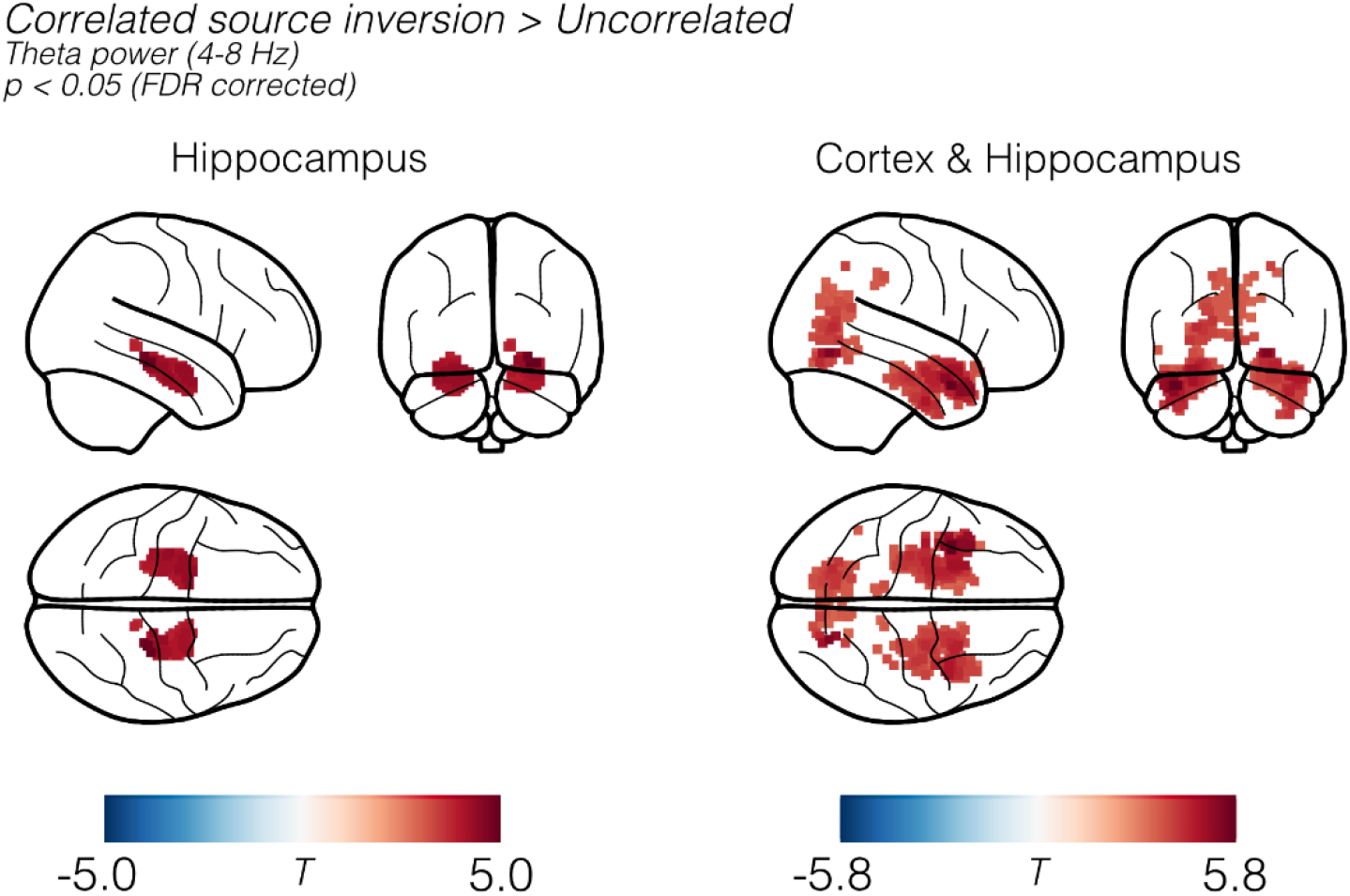
Reconstructed theta band power based on hippocampal only (left panel) and whole-brain (right panel) correlated priors contrasted (paired T-test) against the uncorrelated source model. Left: Comparing uncorrelated to a model containing a correlated hippocampus, we noted a significant increase in theta-band (4-8 Hz) power over the anterior hippocampal areas. Right: Contrasting the whole-brain correlated model with the uncorrelated EBB solution. We observed bilateral anterior hippocampal clusters with significant increases in power in the parahippocampal, rhinal and temporal pole areas. Results have been tranformed into MNI space for visualisation purposes.

## 5) Discussion

We have proposed a method to implement and test between different correlated source models in beamformer neuroimaging. This is possible within an empirical Bayes framework in which we can objectively quantify the likelihood of any particular model given the data. We applied this correlated source framework to experimental MEG data to show, for the first time, that human hippocampal activity in a scene imagination paradigm is bilateral.

We revisited the Empirical Bayesian Beamformer (EBB) in simulation and demonstrated that (unlike normal beamformer solutions) it failed gracefully in the presence of correlated sources. This is because when covariant sources are suppressed, the EBB-prior becomes featureless, approximating a minimum-norm prior. We then introduced the cEBB variant in which the prior variance component consisted of a paired-correlated-source prior plus the standard EBB prior. This meant that the cEBB variant gave qualitatively similar and accurate performance for both correlated and uncorrelated source pairs. Using unilateral, and bilateral correlated and uncorrelated cortical source we showed how model evidence can be used to identify the most likely model for a given dataset. We observed similar performance for source configurations along the body of the hippocampi; however, our inference broke down at the anterior portion of the hippocampi. In this region, although we were able to distinguish between unilateral and bilateral sources, our analysis identified uncorrelated source pairs as correlated. It seems that this was due to the high correlation between the sensor-level profiles produced by the either of the anterior sources (see also Supplementary figure 4) rather than any other factor (such as source distance or dipole orientation). In other words, although spatially distinct, the anterior hippocampal portions occupy the same region in cryogenic MEG signal space. As approximately 50% of the variance in one source will automatically be explained by the other (due to the lead-field correlations), even uncorrelated sources appear as correlated from this viewpoint. It is, however, encouraging that the overlap of sensitivity was not so large that we were unable to distinguish unilateral from bilateral sources.

For the experimental data, we found that adding a bilaterally correlated hippocampal prior gave a consistent improvement in model evidence across the cohort (Fig. 5A). The random effects analysis suggested that this model was not only the most likely, but was clearly distinct from the classical beamformer/EBB solution to the problem (Fig. 5B, 5C). Furthermore, and perhaps more striking, we also observed a significant improvement in model evidence when adding a correlated hippocampus to an already correlated cortex. It is worth noting that in both these winning models, the hippocampal sources only account for 3.9% of the total source space, and these sources were in turn deep in the brain. Making use of the winning correlated-hippocampal model, we noted an increase in theta power in both hippocampi (Fig. 6, left panels) as compared to standard beamformer (EBB) estimates. Also, if we removed the hippocampal specificity and allowed correlation to exist across the whole brain (Fig. 6 right panels), the main power increases were once again observed in these regions. Note, however, that based on our simulations, and the anterior location of the hippocampal power change, we were unable to resolve whether these sources were correlated or not. The conservative conclusion is simply that the power changes are more likely bilateral than unilateral. In the Supplementary Information we investigate what particular properties of the modelled dipoles in the hippocampus might be driving this area of uncertainty on the anterior hippocampus. The results may determine whether potentially adding more anterior or inferior sensors, or even changing the sensor orientation may allow for better separation of these sources. We are currently looking into designs of on-scalp optically-pumped MEG (OP-MEG) systems^58^ that might more effectively distinguish between these anterior hippocampal portions^59^.

It is important to note that the correlated-cortex and whole-brain correlated models generated the largest absolute changes in model evidence (relative to the uncorrelated prior), however in 5/22 participants these changes were in the opposite direction. Based on the random effects analysis, these models were not significantly different from the standard EBB implementation. However, contrasting theta power compared to the EBB model still showed significant increases in power in parahippocampal, rhinal and temporal pole areas (Fig. 6). One explanation is that our anatomical model did not match reality: these data were all source reconstructed using a fitted template anatomical image, with canonical cortical and hippocampal meshes. We know this approach is robust and works well for volumetric studies^15–17,51,60–65^ but it may well be that co-registration errors or individual anatomical variability mean that some hippocampal variance is explained by the cortical mesh or vice versa. Given the large cluster of significant power changes in bilateral temporal lobes, this seems likely. Even when the individual anatomy is available, we know that small errors in co-registration can undermine accurate forward models^66^. The inconsistent improvements/reductions in model evidence could be attributed to forcing 96% of the source space to be correlated in this scenario, when in reality a much smaller area is exhibiting correlated behaviour. As an aside, when using the whole-brain correlated model, we also observed a cluster of theta power located towards the visual cortex, which has been reported in similar paradigms in both MEG^17,67,68^ and fMRI^69^.

The key neuroscience finding from this investigation is that when provoked by an imagination paradigm, theta oscillations within the human hippocampus are bilateral. Our motivation was the discrepancy between functional activation results reported when using fMRI and MEG (with beamformers) to image similar tasks. fMRI experiments designed to engage the hippocampus often report bilateral activation^12,69,70^, whereas there is typically a lateralisation to a single hippocampus in MEG. The experimental MEG data presented in the current study originated from a previous experiment^16^ where the authors found theta band oscillatory activity localised to the left hippocampus and temporal lobe during scene imagination trials. A follow-up study, where scalp-based OP-MEG was compared to conventional SQUID-MEG in the same paradigm showed left lateralised hippocampal theta using SQUID-MEG, but right lateralised hippocampal activity using OP-MEG, despite the same participants being scanned in the two MEG systems^17^. Finally, a recent MEG study investigating autobiographical memory recall reported lateralised activity in the left anterior hippocampus^15^. The common link between these MEG studies is that their source reconstruction was performed using LCMV beamforming, the performance of which is impaired by distal correlated sources, or extended cortical patches^71^ (which is what we effectively have with paired anterior hippocampal sources sharing over 50% variance between them). Indeed, moving away from scene-specific paradigms, and simply counting the number of MEG papers reporting hippocampal activity (see Supplementary Figure 1), we found that 49/83 papers reported unilateral hippocampal activity, and of those that did, 39 used a source reconstruction method within the beamformer family of solutions. Conversely, only 11 beamformer-type reconstructions led to reported bilateral activations in the hippocampus, but even in that case, one of these studies used modified beamformers to specifically image correlated sources^72^.

To summarise, we have explored the possibility that the hippocampus or the cortex or both are correlated in their activity across hemispheres during an imagination paradigm. We found strong evidence that a correlated hippocampal (and uncorrelated cortical) model provided the best explanation of the data. These findings may have implications more generally, by helping to explain why, up until now, the majority of MEG-reported hippocampal activity (typically making use of beamformers) has been estimated as unilateral.

## Acknowledgements

The Wellcome Centre for Human Neuroimaging is supported by a Centre Award from Wellcome (203147/Z/16/Z). Funding for this project derived from the Wellcome Collaborative Award (203257/Z/16/Z) and EPSRC (EP/T001046/1) and the Quantum technology hub in sensing and timing (sub-award QTPRF02). Stephanie Mellor was funded through the EPSRC funded UCL centre for doctoral training (EP/L016478/1) . Eleanor Maguire is supported by a Wellcome Principal Research Fellowship (210567/Z/18/Z).

