## Supplementary information for "Testing covariance models for MEG source reconstruction of hippocampal activity"

### *Supplementary Material*

George C. O'Neill[a], Daniel N. Barry[b], Tim M. Tierney[a], Stephanie Mellor[a], Eleanor A. Maguire[a], Gareth R. Barnes[a]

[a]*Wellcome Centre for Human Neuroimaging, UCL Queen Square Institute of Neurology, University College London, London, UK*
[b]*Department of Experimental Psychology, University College London, London, UK*

---

#### *1) The pattern of hippocampal results in the MEG literature*

To investigate the prevalence of unilateral and bilateral hippocampal activity in reported MEG studies, we performed a miniature review of literature.

##### *Methods*

We searched the PubMed database for all documents that contained both the terms *magnetoencephalography* and *hippocampus* that were published between 1st Jan 1990 and 1st July 2020. We then checked that each paper returned by PubMed met the following criteria:

1. The publication was not a duplicate result of a previously parsed result;
2. The publication was not a literature review or commentary;
3. The publication was not pathological case report;
4. The publication was not a simulation study;
5. The publication contained electrophysiological recordings of humans that covered both hippocampal regions;
6. A source-level analysis of the experimental data was performed;
7. A significant activation or network node was identified in either or both hippocampal regions.

If all criteria were met, we recorded whether there were reported activations in one or both hippocampal regions and identified what type of experiment was performed and what source inversion method was implemented.

##### *Results*

PubMed retuned 191 publications containing both keywords, of which we eliminated 108 for failing to meet all 7 criteria, leaving 83 qualifying publications. Figure S1 is a Sankey diagram of the breakdown of the results. 49/83 (59 %) of publications reported unilateral hippocampal findings with the remaining 34 (41 %) reporting bilateral activity. We also note that the most common source reconstruction method was the beamformer, used in 50/83 (60 %) of studies. Of those 50 beamformer studies, 39 reported unilateral activations in the hippocampus, whilst only 11 noted bilateral activity. L2 minimum norm solutions (which include MNE type and LORETA type source reconstructions) were almost an equal split between unilateral (5) and bilateral (6). Equivalent Current Dipole (ECD) studies predominantly reported bilateral findings (14/18). Finally, both L1 minimum norm solutions reported unilateral activity, and usage of multiple sparse priors (MSP) gave exclusively bilateral findings. The breakdown of which studies reported which results is provided in Table S1

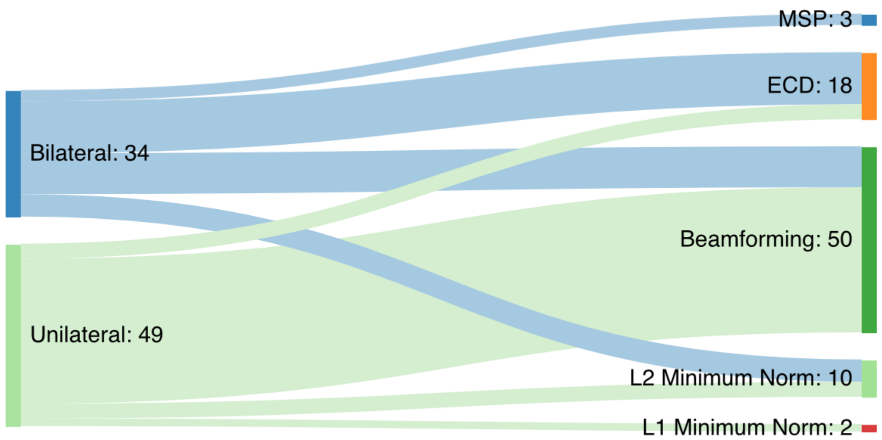


***Figure S1:*** *Sankey flow diagram depicting the split between MEG studies reporting unilateral and bilateral hippocampal activity and which family of inverse solution was implemented.*

| **Inverse Method** | **Hippocampal Response** | **No. Studies** |
|---|---|---|
| Multiple sparse priors (MSP) | Bilateral | 3 [1–3] |
| Equivalent current dipole fitting (ECD) | Bilateral | 14 [4–17] |
| | Unilateral | 4 [18–21] |
| Beamforming | Bilateral | 11 [22–32] |
| | Unilateral | 39 [33–71] |
| L2 Minimum Norm | Bilateral | 6 [72–77] |
| | Unilateral | 4 [78–81] |
| L1 Minimum Norm | Unilateral | 2 [82,83] |

***Table S1:*** *Tabular breakdown of the results.*

***2) The effect of not accounting for uncorrelated source variance in prior selection***

Within the main manuscript, we summed the variance between uncorrelated and correlated sources when designing our priors for cEBB inversion. Whether this is the most appropriate approach is an open question, but here we characterise what would happen if uncorrelated source variance is not accounted for.

Recall Equation 7 in the main manuscript, where we defined the cEBB source covariance matrix as the sum of the original EBB prior and the variance from a set of correlated sources $\boldsymbol{p}'$.

$$\boldsymbol{Q}_{j|cEBB} = \boldsymbol{Q}_{j|EBB} + \text{diag}(\boldsymbol{p}'^2). \qquad \text{(S1)}$$

For this demonstration we shall define an exclusively correlated EBB (xcEBB) covariance matrix as

$$\boldsymbol{Q}_{j|xcEBB} = \text{diag}(\boldsymbol{p}'^2). \qquad \text{(S2)}$$

For simplicity, we compared cEBB and xcEBB on our simulations within Heschl's gyri.

*Results*

Figure S2 shows the changes in model evidence where comparing EBB to either cEBB or xcEBB, we see that the two variations of correlated source inversions agreed with each other when selecting the most plausible model. For single sources and uncorrelated, an uncorrelated model (EBB) won, whereas for correlated sources both the correlated models (cEBB and xcEBB) up until SNR values below -30 dB, where it breaks down and believed the correlated sources are in fact not. We observed that the magnitude of the changes were larger for xcEBB models than cEBB models. Figure S3A shows the spatial distribution of the source variance priors used in the model inversions for the mono simulations. Two points are noteworthy, the variance projected into the contralateral hemisphere of the original source relative to the variance in the ipsilateral hemisphere was higher in the xcEBB model than compared to cEBB, a consequence of the variance from the uncorrelated sources being absent. We also noted that in the ipsilateral hemisphere the variance in xcEBB was projected to be more superficial than for cEBB, which was reflected in the localisation of the power, as shown in Figure S3B.

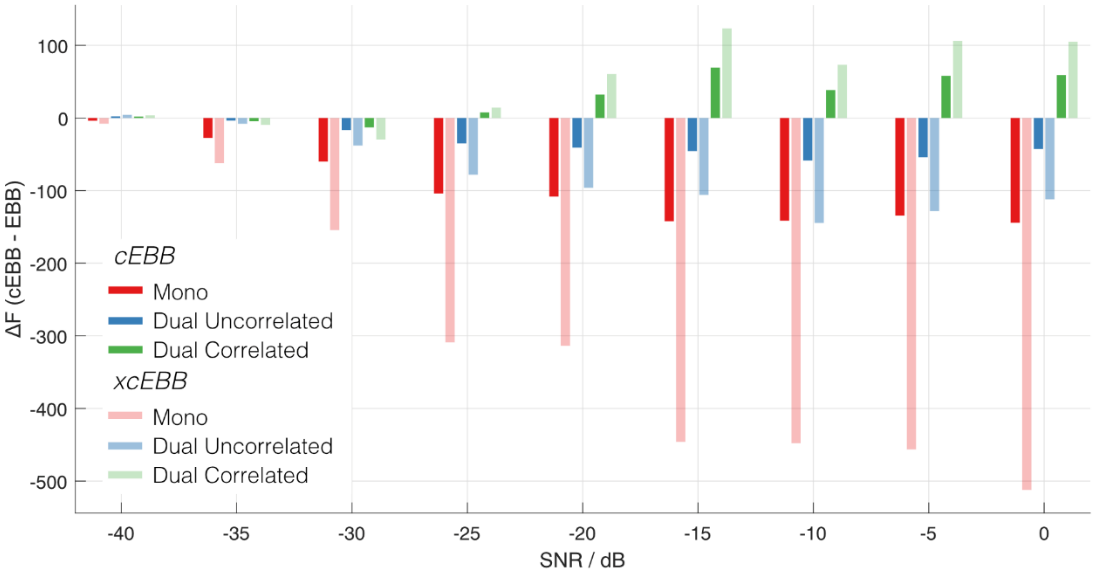


***Figure S2:*** *Model evidence changes when comparing correlated source priors compared EBB for sources within Heschl's gyri. Darker bars represent cEBB where the prior consists of a mix of correlated and uncorrelated source variance, lighter colours represent xcEBB which is purely made from correlated sources.*

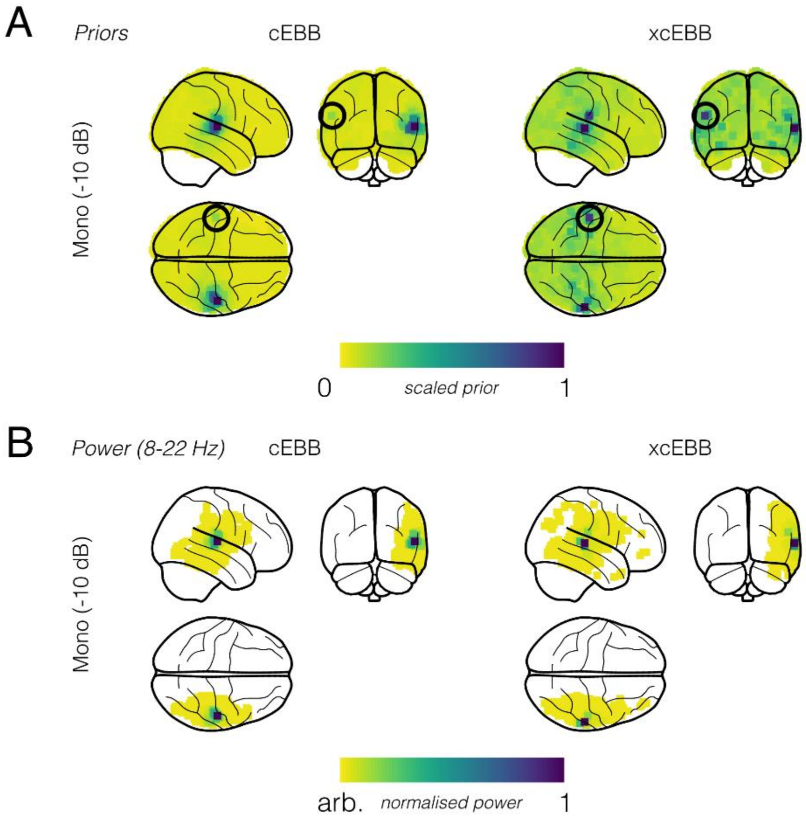


***Figure S3:*** *A comparison of cEBB and xcEBB in a single source simulation in Heschl's gyri. A) Spatial distribution of the priors. The solid black circle represents where the correlated source assumptions tried to project a single source into the other hemisphere. xcEBB did this to a larger extent than cEBB. B) Source reconstructed power of oscillations between 8-22 Hz.*

#### *3) The relationships between dipole properties and model evidence*

Within the main manuscript we speculated that the difficulty in separating bilateral anterior hippocampal sources is largely attributed to the high correlation of the lead fields between the two areas. Here we probe whether some other properties of the sources may also contribute.

*Methods*

We repeated the dual uncorrelated source simulations from the main manuscript (SNR fixed to -10 dB), but over 256 randomly sampled homologous source pairs in the cortex and 163 hippocampal homolog pairs. We compared the changes in model evidence between the EBB and cEBB model inversions. We would expect the introduction of a correlated source prior to see a reduction in the model evidence. In particular we focussed on 4 properties of the sources:

- The correlation of the lead fields between the source pairs
- The 2-norm of the lead fields
- The distance between the source pairs
- The orientation of the sources relative to the radial direction of a single sphere fitted to the anatomy.

*Results*

Figure S4 shows the scatter plots relations between switching source models and the properties of the bilateral dipole sources, cortical sources in grey and hippocampal sources in dark blue for contrast. We see a clear relationship between the correlations in the sources lead fields and the change in model evidence (Fig. S4A). Interestingly, we observed that if the two source lead fields were strongly anticorrelated then the correct model EBB (as represented as a reduction in model evidence when switching to cEBB) was selected for these simulations. But if the lead fields were positively correlated, cEBB was regarded as the more plausible solution. We observed that in the hippocampus there was a trend for closer sources to prefer the cEBB model, something that we did not observe in the cortex (Fig 4C). The lead field norms (Fig. 4B) and dipole orientation (Fig. 4D) did not show any obvious preference.

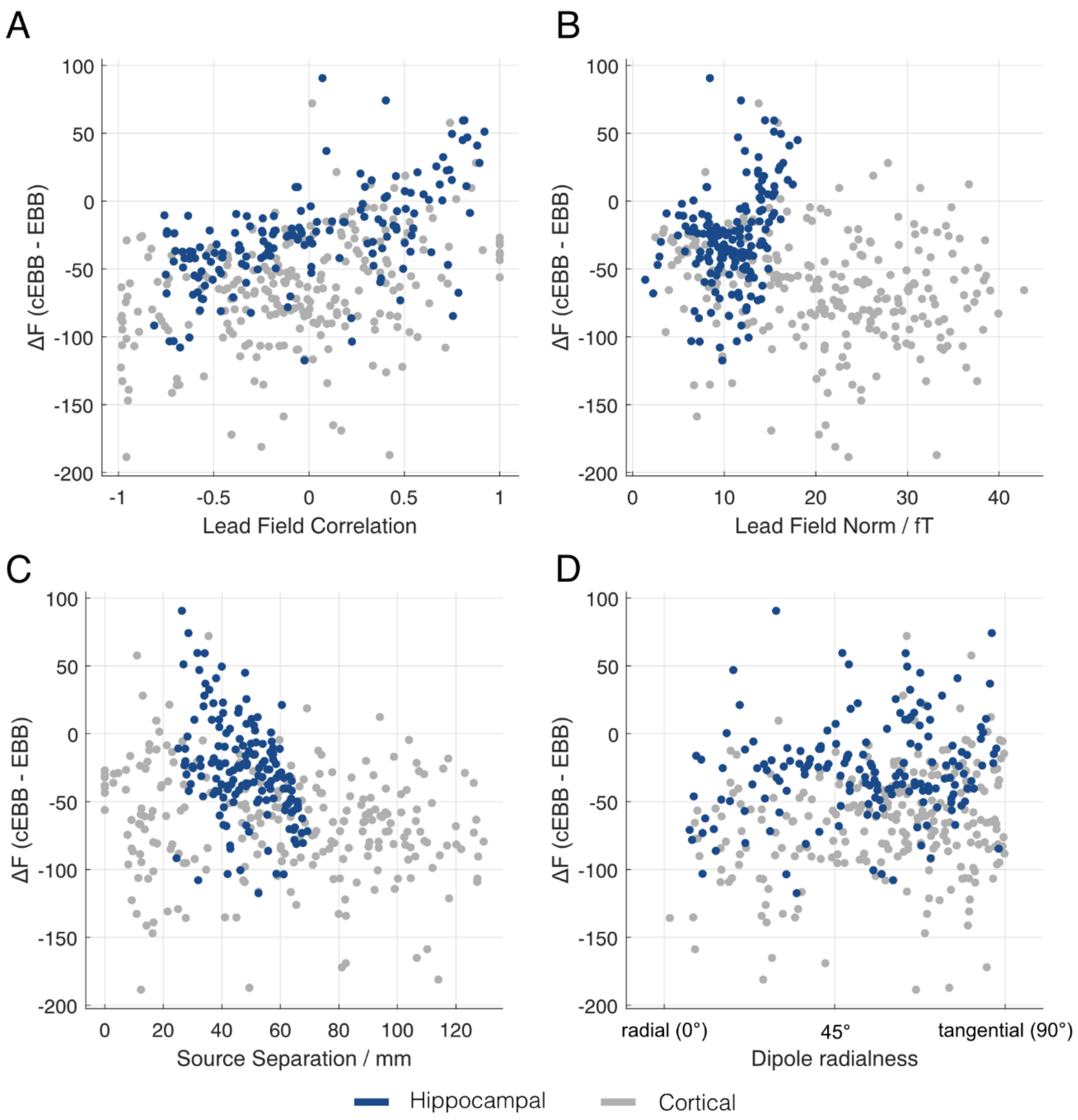


***Figure S4:*** *Scatter plots showing relationships between changes in model evidence and various dipole properties within the hippocampus (dark blue dots) and cortex (grey dots).*

### *Bibliography*


1. Urbain, C. *et al.* Sleep in children triggers rapid reorganization of memory-related brain processes. *Neuroimage* **134**, 213–222 (2016).
2. Tzovara, A. *et al.* High-precision magnetoencephalography for reconstructing amygdalar and hippocampal oscillations during prediction of safety and threat. *Hum. Brain Mapp.* **40**, 4114–4129 (2019).
3. Campo, P. *et al.* Remote Effects of Hippocampal Sclerosis on Effective Connectivity during Working Memory Encoding: A Case of Connectional Diaschisis? *Cereb. Cortex* **22**, 1225–1236 (2012).
4. Nishitani, N. The role of the hippocampus in auditory processing studied by event-related electric potentials and magnetic fields in epilepsy patients before and after temporal lobectomy. *Brain* **122**, 687–707 (1999).
5. Nishitani, N. Dynamics of cognitive processing in the human hippocampus by neuromagnetic and neurochemical assessments. *Neuroimage* **20**, 561–571 (2003).
6. Leirer, V. M. *et al.* Hippocampal activity during the transverse patterning task declines with cognitive competence but not with age. *BMC Neurosci.* **11**, 113 (2010).
7. Kikuchi, Y. *et al.* Human cortico-hippocampal activity related to auditory discrimination revealed by neuromagmetic field. *Neuroreport* **8**, 1657–1661 (1997).
8. Williams, T. J., Nuechterlein, K. H., Subotnik, K. L. & Yee, C. M. Distinct neural generators of sensory gating in schizophrenia. *Psychophysiology* **48**, 470–478 (2011).
9. Hamada, Y., Sugino, K., Kado, H. & Suzuki, R. Magnetic fields in the human hippocampal area evoked by a somatosensory oddball task. *Hippocampus* **14**, 426–433 (2004).
10. Papanicolaou, A. C. The Hippocampus and Memory of Verbal and Pictorial Material. *Learn. Mem.* **9**, 99–104 (2002).
11. Kirsch, P. *et al.* Cerebellar and Hippocampal Activation During Eyeblink Conditioning Depends on the Experimental Paradigm: A MEG Study. *Neural Plast.* **10**, 291–301 (2003).
12. Hanlon, F. M. *et al.* A Specific Test of Hippocampal Deficit in Schizophrenia. *Behav. Neurosci.* **119**, 863–875 (2005).
13. Tesche, C. D. & Karhu, J. Theta oscillations index human hippocampal activation during a working memory task. *Proc. Natl. Acad. Sci.* **97**, 919–924 (2000).
14. Nishitani, N., Nagamine, T., Fujiwara, N., Yazawa, S. & Shibasaki, H. Cortical-Hippocampal Auditory Processing Identified by Magnetoencephalography. *J. Cogn. Neurosci.* **10**, 231–247 (1998).
15. Maestú, F. *et al.* Biomagnetic profiles of verbal memory success in patients with mesial temporal lobe epilepsy. *Epilepsy Behav.* **16**, 527–533 (2009).
16. Tesche, C. D., Karhu, J. & Tissari, S. O. Non-invasive detection of neuronal population activity in human hippocampus. *Cogn. Brain Res.* **4**, 39–47 (1996).
17. Hanlon, F. M. *et al.* A non-invasive method for observing hippocampal function. *Neuroreport* **14**, 1957–1960 (2003).
18. Ueda, Y. *et al.* The presence of short and sharp MEG spikes implies focal cortical dysplasia. *Epilepsy Res.* **114**, 141–146 (2015).
19. Ver Hoef, L. W., Sawrie, S., Killen, J. & Knowlton, R. C. Left Mesial Temporal Sclerosis and Verbal Memory: A Magnetoencephalography Study. *J. Clin. Neurophysiol.* **25**, 1–6 (2008).
20. Breier, J. I., Simos, P. G., Zouridakis, G. & Papanicolaou, A. C. Lateralization of cerebral activation in auditory verbal and non-verbal memory tasks using magnetoencephalography. *Brain Topogr.* **12**, (1999).
21. Tesche, C. D. & Karhu, J. Interactive Processing of Sensory Input and Motor Output in the Human Hippocampus. *J. Cogn. Neurosci.* **11**, 424–436 (1999).
22. Mišić, B. *et al.* Post-Traumatic Stress Constrains the Dynamic Repertoire of Neural Activity. *J. Neurosci.* **36**, 419–431 (2016).
23. Costers, L. *et al.* Spatiotemporal and spectral dynamics of multi-item working memory as revealed by the n -back task using MEG. *Hum. Brain Mapp.* **41**, 2431–2446 (2020).
24. Rondina, R. *et al.* Age-related changes to oscillatory dynamics in hippocampal and neocortical networks.

*Neurobiol. Learn. Mem.* **134**, 15–30 (2016).
